# Process Development and Scale-up Optimization of the SARS-CoV-2 Receptor Binding Domain-Based Vaccine Candidate, RBD219-N1C1

**DOI:** 10.1101/2020.12.30.424829

**Authors:** Jungsoon Lee, Zhuyun Liu, Wen-Hsiang Chen, Junfei Wei, Rakhi Kundu, Rakesh Adhikari, Joanne Altieri Rivera, Portia M. Gillespie, Ulrich Strych, Bin Zhan, Peter J. Hotez, Maria Elena Bottazzi

## Abstract

A SARS-CoV-2 RBD219-N1C1 (RBD219-N1C1) recombinant protein antigen formulated on Alhydrogel^®^ has recently been shown to elicit a robust neutralizing antibody response against SARS-CoV-2 pseudovirus in mice. The antigen has been produced under current good manufacturing practices (cGMP) and is now in clinical testing. Here, we report on process development and scale-up optimization for upstream fermentation and downstream purification of the antigen. This includes production at the 1 and 5 L scale in the yeast, *Pichia pastoris,* and the comparison of three different chromatographic purification methods. This culminated in the selection of a process to produce RBD219-N1C1 with a yield of >400 mg per liter of fermentation with >92% purity and >39% target product recovery after purification. In addition, we show the results from analytical studies, including SEC-HPLC, DLS, and an ACE2 receptor binding assay that were performed to characterize the purified proteins to select the best purification process. Finally, we propose an optimized upstream fermentation and downstream purification process that generates quality RBD219-N1C1 protein antigen and is fully scalable at a low cost.

## Introduction

After the first report of coronavirus disease 2019 (COVID-19) in December 2019 [21,24], the number of cases is now at 75 million with over 1.6 million deaths worldwide [16]. As of December 2020, two vaccines from Pfizer/BioNTech and Moderna have been approved for emergency use in multiple countries. These frontrunner vaccines represent a new platform, mRNA-based vaccines; while they were produced in record time, they are relatively expensive to manufacture and require transportation and storage at temperatures below −20°C. Such features present formidable challenges to distribute these vaccines to low- and middle-income countries (LMICs). Confounding equity access for COVID-19 vaccines are the still uncertain production and distribution delays or even the quality of other vaccines. This situation could leave LMICs bereft of low-cost COVID-19 vaccines suitable for their modest or depleted health systems [20]. In response, a network of LMIC vaccine developers and manufacturers are accelerating vaccines employing traditional platforms, such as an inactivated virus and recombinant proteins, with several under preclinical and clinical development [9]. These vaccine candidates are less demanding with respect to transport and storage, and often come with a long history of successful production and use for other infectious diseases [15]. Particularly attractive in this aspect are recombinant protein antigens produced through microbial fermentation in yeast. For instance, recombinant Hepatitis B vaccine has been administered to adults and children for decades [13].

The SARS-CoV-2 virus, the pathogen that causes COVID-19, uses its surface spike (S) protein for host cell entry, just like its close relative, SARS-CoV that had caused an outbreak of Severe Acute Respiratory Disease in 2002. The Receptor Binding Domain (RBD) of the S protein binds to a cellular receptor, angiotensin-converting enzyme 2 (ACE2) that mediates membrane fusion during viral entry into the cell [22,28]. S proteins for both viruses have served as vaccine antigens that could elicit antibodies to prevent virus entry by blocking the binding of RBD to ACE2 [14], and all the leading COVID-19 vaccines currently in clinical trials, including the mRNA vaccines, use the S protein to elicit immunity [12]. Overwhelmingly, such vaccines protect through their induction of virus-neutralizing antibodies, together with T cell responses [17].

Building on our experience with the RBD of the SARS-CoV spike protein [5,7,4], the SARS-CoV-2 RBD was cloned and expressed in the yeast *Pichia pastoris* [8]. Yeast has a track record of serving as a host organism for the production of multiple regulatory-approved and prequalified recombinant subunit vaccines, including vaccines for Hepatitis B, Influenza B, human papillomavirus, as well as for Diphtheria and Tetanus [1,19]. Eukaryotic expression in yeast shows advantages over the prokaryote, *Escherichia coli*, with respect to the production of recombinant protein vaccines. Proper protein folding, disulfide bridge formation, post-translational modifications, and secretory cleavage are better supported in yeast, while also allowing for robust production with low costs and full scalability, features that distinguish this platform from other eukaryotic systems, such as insect cells, mammalian cells, and plants.

The RBD219-N1C1 antigen is derived from residues 332-549 of the SARS-CoV-2 RBD with a single mutation of a free cysteine residue (Cys538) to alanine to prevent intermolecular disulfide bond formation and therefore unwanted oligomerization during process development [8]. In addition, N1 refers to the deletion of Asn331 to avoid hyperglycosylation observed in previous studies with the SARS-CoV RBD219-N1 antigen [5]. In initial studies, the RBD219-N1C1 recombinant protein antigen adjuvanted with Alhydrogel^®^ has been shown to elicit a robust neutralizing antibody response against SARS-CoV-2 pseudovirus in mice [25].

With any COVID-19 vaccine candidate, the ability to produce billions of doses efficiently is crucial to satisfy the potential global vaccine demand. We, therefore, have been developing and optimizing a scalable production process of the RBD219-N1C1 vaccine candidate at low cost to support its technology transfer. Initial fermentation runs scouting for growth media, induction time, and glycerol fed-batch conditions were executed in a 1 L bioreactor and resulted in a ~10-fold increase in RBD219-N1C1 expression levels. Further scale-up experiments in a 5 L bioreactor established reproducibility of the selected conditions. Simultaneously, a purification scheme was developed based on the process used for the 70% homologous SARS-CoV RBD antigen [4], and further optimized to allow full scalability and lower the cost. Taking into consideration, yield, purity, functionality, and removal of host cell contaminants, we have developed an optimized fermentation at the 5 L scale and purification *(Process-2)* suitable for production and manufacturing of a high-yield (and therefore potentially low-cost) COVID-19 vaccine antigen candidate. The developed process has already been transferred to an industrial vaccine manufacturer in India and is currently undergoing further production maturity while the vaccine candidate is in a Phase 1/2 clinical trial.

## Materials and Methods

### Generation of Research Cell Bank

To generate a research cell bank (RCB), *P. pastoris* X33 strain was transformed with expression plasmid pPICZαA containing RBD219-N1C1 coding DNA, and one transformed colony with high expression of recombinant RBD219-N1C1 protein [8] was selected and streaked on YPD plates containing 100 μg/mL Zeocin to make single colonies. The plates were incubated at 30 °C for approximately 3 days until single colonies were observed. Subsequently, 200 mL plant-derived phytone YPD medium was inoculated with a single colony from the respective plate and incubated at 30 °C with constant shaking (225 RPM) until the OD_600_ reached 9.3. Finally, the cell culture was mixed with plant-derived glycerol to a final concentration of 20% and aseptically aliquoted (1 mL each) into 1.2 mL cryovials. For longterm storage, the cryovials were stored at −80 °C.

### Fermentation

One vial of the SARS-CoV-2 RBD219-N1C1 RCB was used to inoculate 1 L BMG (Buffered Minimal Glycerol) medium in a 2 L shake flask. The shake flask culture was grown at 30°C and 225 rpm until an OD_600_ of 5 – 10. For 1 L fermentations, this seed culture was inoculated into 0.4 L of sterile basal-salt medium (BSM), pH 5.0 (BSM: 18.2 g/L potassium sulfate, 14.9 g/L magnesium sulfate heptahydrate, 4.13 g/L potassium hydroxide, 0.93 g/L calcium sulfate dehydrate, 26.7 mL/L of 85% phosphoric acid, and 40 g/L glycerol) or low-salt medium (LSM), pH 5.0 (LSM: 4.55 g/L potassium sulfate, 3.73 g/L magnesium sulfate heptahydrate, 1.03 g/L potassium hydroxide, 0.23 g/L calcium sulfate dehydrate, 10.9 mL/L of 85% phosphoric, and 40 g/L glycerol) to a starting cell density (OD_600_) of 0.5. Fermentation was conducted using a Biostat Qplus Bioreactor with a 1 L vessel (Sartorius Stedim, Guxhagen, Germany). For 5 L runs, the seed culture was inoculated into 2.5 L of LSM, and fermentation was conducted in a CelliGen 310 Bioreactor with a 7.5 L vessel (Eppendorf, New York, USA), controlled by the Eppendorf Bio Command software. Cell expansion was continued at 30 °C with a dissolved oxygen (DO) set point of 30%. Following the DO spike, a fed-batch was initiated with 50% glycerol at a feed rate of 15 mL/L/hr for 6 hours. During the last hour of the fed-batch phase, pH and temperature were increased to the desired value (pH=6.5, temperature=25°C). When a glycerol fed-batch was not included in the fermentation process, the pH and temperature were adjusted to the desired value during the first hour of induction. After the fed-batch phase, methanol induction was initiated; the total induction time was approximately 68-72 hours. Biomass was removed by centrifugation at 12,227 x g for 30 minutes at 4 °C before the supernatant was filtered through 0.45 μm PES filters stored at −80 °C until purification.

### Purification overview of three processes

The FS was removed from −80 °C and thawed at 22 °C for 4-6 hours. Three purification processes were performed with 1 L FS aliquots (Figure 1B). In Process-1, the RBD219-N1C1 protein was captured from the FS using hydrophobic interaction chromatography (HIC), concentrated by ultrafiltration/diafiltration (UFDF), and polished using size exclusion chromatography (SEC). In Process-2, the RBD219-N1C1 protein was captured using HIC, buffer-exchanged (UFDF), and polished using anion exchange chromatography (AEX). Finally, in Process-3, the FS was buffer-exchanged using UFDF before the target protein was captured using cation exchange chromatography (CEX), buffer-exchanged (UFDF), and polished using AEX.

**Figure 1.**
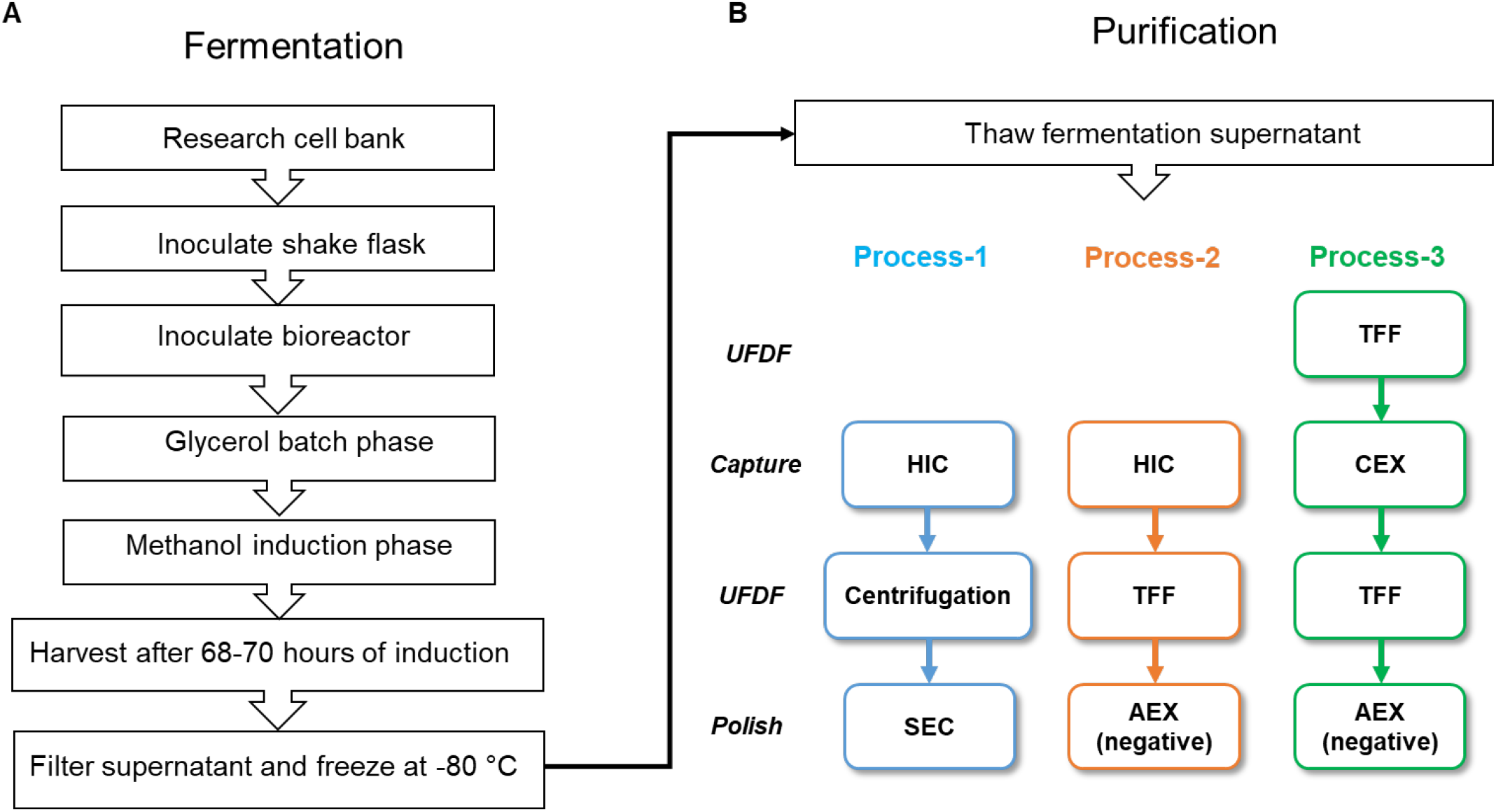
Fermentation flow diagram (A) and purification flow diagram (B). Three purification processes performed are shown in different colors. The color scheme remains consistent throughout all figures. UFDF, Ultrafiltration, and diafiltration; HIC, hydrophobic interaction chromatography; SEC, size exclusion chromatography; TFF, tangential flow filtration; CEX, cation exchange chromatography; AEX, anion exchange chromatography

### UFDF (Ultrafiltration and Diafiltration)

Two types of devices were used for UFDF, a centrifugal concentrator, and a flat sheet membrane, depending on the target volume. For Process-1, Amicon centrifugal concentrator, with a 10 kDa molecular weight cutoff (MWCO) (MilliporeSigma, Burlington, USA) was used to concentrate the HIC elution pool (2,050 x g at 4°C). This allowed concentration to the small volume needed for SEC. For Process-2, a flat sheet Pellicon XL Cassette with a Biomax-5 membrane (5 kDa MWCO) and a LabScale TFF System (MilliporeSigma, Burlington, USA) were used to concentrate the HIC elution pool 8-fold, followed by diafiltration with 4 diavolumes of 20 mM Tris-HCl pH 7.5 and 100 mM NaCl. A crossflow was kept at 25 mL/min over a 0.005 m^2^ membrane area throughout the entire process with an average transmembrane pressure (TMP) of ~15 psi. For Process-3, a flat sheet Pellicon 2 Mini Cassette with a Biomax-5 membrane (MilliporeSigma, Burlington, USA) was used for the first UFDF (UFDF-1) to concentrate the FS 4-fold followed by diafiltration with 4 diavolumes of 20 mM sodium citrate pH 4.2 and 10 mM NaCl. A crossflow was kept constant at 200 mL/min over a 0.1 m^2^ membrane area throughout the entire process with an average TMP of ~8 psi. For the UFDF-2, the CEX elution pool was concentrated 4-fold followed by diafiltration with 4 diavolumes of 25 mM Tris-HCl pH 7.2 and 5 mM NaCl using the Pellicon XL Cassette as described for Process-2.

### Hydrophobic interaction chromatography (HIC)

In Processes 1 and 2, HIC was used to capture RBD219-N1C1 proteins from the FS. Ammonium sulfate salt was added to the FS to a final concentration of 1 M (w/v) and the pH was adjusted to 8.0. The FS was filtered through a 0.45 μm PES filter unit and loaded on a 112 mL Butyl Sepharose High-Performance column (4.4 cm diameter and 7.4 cm bed height) at 20 mL/min flow rate. The column was washed with 1 M ammonium sulfate in 30 mM Tris-HCl pH 8.0. Bound proteins were eluted with 0.4 M ammonium sulfate in 30 mM Tris-HCl pH 8.0.

### Size exclusion chromatography (SEC)

Five mL of the concentrated HIC elution pool was loaded on a HiLoad 16/600 Superdex 75 prep-grade column (Cytiva, Marlborough, USA), pre-equilibrated with 20 mM Tris-HCl pH 7.5 and 150 mM NaCl, and eluted at a flow rate of 1 mL/min. The SEC elution pool was aseptically filtered using 0.2 μm PES in a biosafety cabinet and stored at −80 °C until usage.

### Ion exchange chromatography (IEX)

In Process-3, RBD219-N1C1 was captured using CEX. The Pellicon 2 retentate pool in 20 mM sodium citrate pH 4.2 and 10 mM NaCl was loaded on a 50 mL CM Sepharose Fast Flow column (2.6 cm diameter and 9.3 cm bed height) at 10 mL/min flow rate. The column was washed with 20 mM sodium citrate pH 4.2 and 10 mM NaCl. Bound proteins were eluted in 20 mM sodium citrate pH 6.6 and 10 mM NaCl.

In Processes 2 and 3, RBD219-N1C1 was polished using a negative capture AEX. The Pellicon XL retentate pool was loaded on a HiPrep Q Sepharose XL 16/10 column (Cytiva, Marlborough, USA) that was pre-equilibrated with 20 mM Tris-HCl pH 7.5 and 100 mM NaCl for Process-2, and 25 mM Tris-HCl pH 7.2 and 5 mM NaCl for Process-3. The flowthrough from AEX was collected, aseptically filtered using 0.2 μm PES filters in a biosafety cabinet, and stored at −80 °C until usage. NaCl (95 mM) was added to the final purified proteins from Process-3 prior to storage in 25 mM Tris-HCl pH 7.2 and 100 mM NaCl.

### Protein yield and purity determination by quantitative SDS-PAGE

In-process samples taken at each purification step were loaded on either 14% Tris-glycine gels or 4-12% Bis-Tris gels to determine the concentration and purity of the various RBD219-N1C1 samples. Purified RBD219-N1C1 proteins of known concentrations were used as standards. After SDS-PAGE, gels were stained with Coomassie Blue and scanned with a GS-900 densitometer (Bio-Rad, Hercules, USA). Gel images were processed with ImageLab software (Bio-Rad, Hercules, USA) to create a standard curve and determine protein concentration and purity.

### Western Blot

Western Blot analysis was performed to detect RBD219-N1C1 as well as *P. pastoris* host cell protein (HCP). Five micrograms of purified protein were run on 14% Tris-Glycine gels under non-reducing and reducing conditions to detect RBD219-N1C1 and HCP, respectively. Proteins in gel were transferred to PVDF membranes and blocked with 5% dry milk in PBST (1X PBS with 0.05% Tween-20). RBD219-N1C1 was detected using a rabbit monoclonal antibody against the SARS-CoV-2 Spike S1 protein (Sino Biological, Beijing, China; Cat#: 40150-R007) and goat anti-rabbit IgG secondary antibodies conjugated with horseradish peroxidase (Invitrogen, Carlsbad, USA; Cat#: G21234). HCPs were detected using an anti-*P. pastoris*:HRP conjugate (2G) solution (Cygnus, Southport, USA; Cat# F641-12). The blots were developed using ECL Prime Substrate System (Cytiva, Marlborough, USA).

### Size Exclusion Chromatography-High Performance Liquid Chromatography (SEC-HPLC)

Waters^^®^^ Alliance HPLC Separations Modules and Associated PDA Detectors were operated to analyze the size and purity of purified RBD219-N1C1 proteins. Fifty μg micrograms of the RBD219-N1C1 protein were injected into a Yarra SEC-3000 column (300 mm X 7.8 mm; catalog # 00H-4513-K0), and was eluted in 20 mM Tris-HCl pH 7.5 and 150 mM NaCl, at the flow rate of 0.6 mL/min. The elution of protein was confirmed by detecting the absorbance at 280 nm.

### Dynamic Light Scattering (DLS)

The size of the purified RBD219-N1C1 proteins was also analyzed by DLS [4,6]. In short, RBD219-N1C1 was first diluted to 1 mg/mL with TBS, and approximately 40 μL of protein were then loaded into a clear bottom 384-well plate in four replicates to evaluate the hydrodynamic radius and molecular weight using the cumulant fitting on a Wyatt Technology DynaPro Plate Reader II.

### Host Cell Protein Quantification by ELISA

Yeast-expressed RBD219-N1C1 is N-glycosylated [8]. To avoid any cross-reactivity from anti-*P. pastoris* HCP antibodies that recognize the N-glycans, which could result in an overestimation of true HCP, we performed quantitative ELISAs with a second-generation anti-*Pichia pastoris* HCP ELISA Kit (Cygnus, Southport, USA; Cat# F640) following the manufacturer’s instructions. This kit provides strips pre-coated with anti-*P. pastoris* HCP antibodies. Serially-diluted RBD219-N1C1 was loaded onto the strips (HCP standards range from 0-250 ng/mL) in the presence of HRP conjugated anti-*P. pastoris* antibodies. The strips were then incubated for approximately 3 hours at room temperature followed by 4 washes. Finally, 100 μL of TMB solution were added to react with the HRP conjugated antibodies that were presented in the strip for 30 minutes prior to the addition of 100 μL of 1 M HCl to stop the reaction. The absorbance of 450 nm was measured in each well of the strip and a linear standard curve was generated by plotting an “absorbance *vs* concentration” graph with the HCP standards to further calculate the HCP concentration present in the RBD219-N1C1 proteins.

### Endotoxin test

Endotoxin levels in the purified RBD219-N1C1 samples were measured using the Endosafe Portable Testing System (Charles River Laboratory, Wilmington, USA). The purified protein was diluted 10-fold with Endosafe LAL reagent water and 25 μL of diluted protein was loaded to each of the four wells of PTS20 Limulus amebocyte lysate Reagent Cartridge for the measurement as described in the literature (Charles River Laboratory, Wilmington, USA) [18].

### *In vitro* ACE2 binding ELISA

The binding of RBD219-N1C1 to recombinant human angiotensin-converting enzyme 2 (ACE2) was evaluated using an ELISA procedure described previously [8]. In short, 96-well ELISA plates were coated with 100 μL 2 μg/mL RBD219-N1C1 overnight at 4°C followed by blocking with PBST/0.1% BSA. 100 μL serially-diluted ACE2-hFc (LakePharma, San Carlos, USA; Cat # 46672) was added to the wells and incubated at room temperature for 2 hours and the binding was detected by adding 100 μL 1:10,000 diluted HRP conjugated anti-human IgG antibodies (GenScript, Piscataway, USA; Cat# A00166) with a 1-hour incubation period at room temperature. Finally, 100 μL TMB substrate was provided to each well to react with HRP and the reaction was terminated with 100 μL 1M HCl. Absorbance at 450 nm was measured using an EPOCH 2 microplate reader (BioTek, Winooski, USA).

## Results

### Fermentation optimization

When basal-salt medium (BSM) and low-salt medium (LSM) were compared for the production of the RBD219-N1C1 protein, no differences were observed in the growth profiles and the final biomass. However, the salt concentration appeared to have a significant effect on the yield. The yield of the RBD219-N1C1 protein using BSM was only 52 mg/L while using LSM 237 mg/L were achieved (Table 1, Runs 1 and 2). Therefore, LSM was used for the further development of the fermentation process.

**Table 1.**
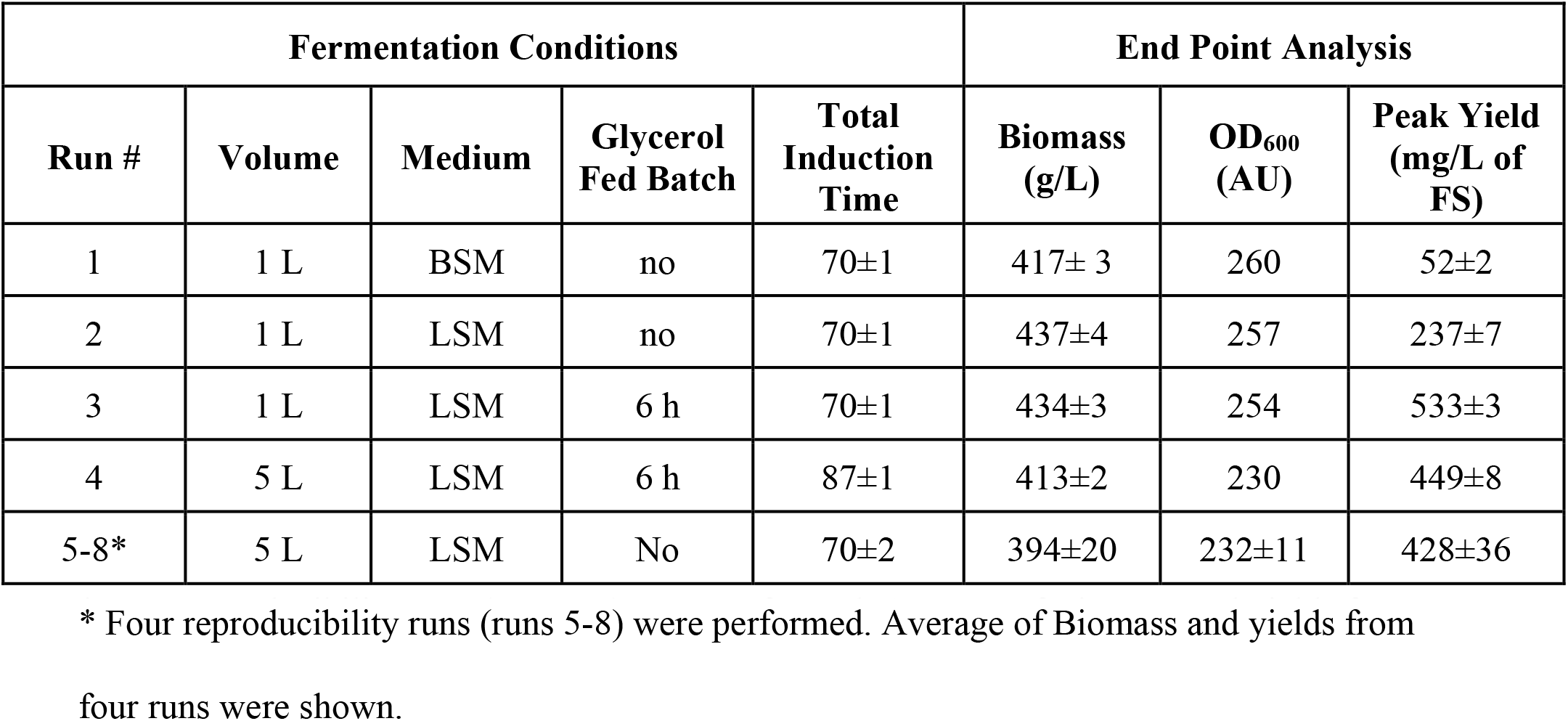
Summary of the development fermentation runs. FS: fermentation supernatant * Four reproducibility runs (runs 5-8) were performed. Average of Biomass and yields from four runs were shown.

| Fermentation Conditions |  |  |  |  | End Point Analysis |  |  |
| --- | --- | --- | --- | --- | --- | --- | --- |
| Run # | Volume | Medium | Glycerol Fed Batch | Total Induction Time | Biomass (g/L) | OD<sub>600</sub> (AU) | Peak Yield (mg/L of FS) |
| 1 | 1 L | BSM | no | 70±1 | 417± 3 | 260 | 52±2 |
| 2 | 1 L | LSM | no | 70±1 | 437±4 | 257 | 237±7 |
| 3 | 1 L | LSM | 6 h | 70±1 | 434±3 | 254 | 533±3 |
| 4 | 5 L | LSM | 6 h | 87±1 | 413±2 | 230 | 449±8 |
| 5-8* | 5 L | LSM | No | 70±2 | 394±20 | 232±11 | 428±36 |

The baseline fermentation process consisted of two phases: a glycerol-batch phase and a methanol fed-batch phase. In glycerol-batch mode, LSM contained 40 g/L of glycerol. At the time of glycerol depletion, the initial induction biomass was 110 ± 10 g/L (WCW). In this study, a glycerol fed-batch phase was then added before methanol induction to test the efficiency of protein expression based on the initial induction biomass. After a 6-hour glycerol fed-batch phase, the initial induction biomass doubled to 210 ± 20 g/L (WCW). The methanol feed strategies were kept the same. At harvest, the final OD_600_ and the biomass were determined to be 260 AU and 417 g/L, respectively. By adding the glycerol fed-batch, the yield of RBD219-N1C1 was increased about 120% to 533 mg/L (Table 1, Run 3).

### Fermentation scalability and reproducibility

The fermentation process with six hours of glycerol feed was then scaled up from 1 L to 5 L to test scalability and reproducibility (Table 1, Run 4). The induction time was extended to 87 hours until biomass started to drop. This suggested that the cells were no longer actively dividing. Since excessive methanol feeding may lead to cell death thus leading to a loss of protein yield, it was decided to stop the methanol feeding after 87 hours of induction. The peak yield of RBD219-N1C1 was 449 ± 8 mg/L at 70 hours after induction (Day 3), after which the yield slightly dropped to 408 ± 9 mg/L at 87 hours after induction.

The fermentation process without the 6-hour glycerol fed-batch phase was also scaled up from 1 to 5 L for comparison (Table 1, Run 5). After 70 ± 2 hours of induction, the yield of RBD219-N1C1 reached 479 ± 15 mg/L (a 128% increase compared to the 1 L scale). This yield was close to the yield of the fermentation run with the 6-hour glycerol fed-batch phase (Table 1, Run 4). Since there was no significant increase in yield by the glycerol fed-batch at 5 L scale, we decided to proceed without this step (Figure 1A). To establish reproducibility, this fermentation process (Figure 1A) was repeated four times (Runs 5-8). The average yield of four reproducibility runs was 428 ± 36 mg/L, with a coefficient of variance of 8.3%. The SDA-PAGE gels analysis of fermentation supernatants of a representative run (Run 5) with the lockdown process was shown in Figure 2. RBD219-N1C1 (a dominant protein band of ~28 kDa) was secreted and accumulated in the fermentation supernatant through the course of methanol induction.

**Figure 2.**
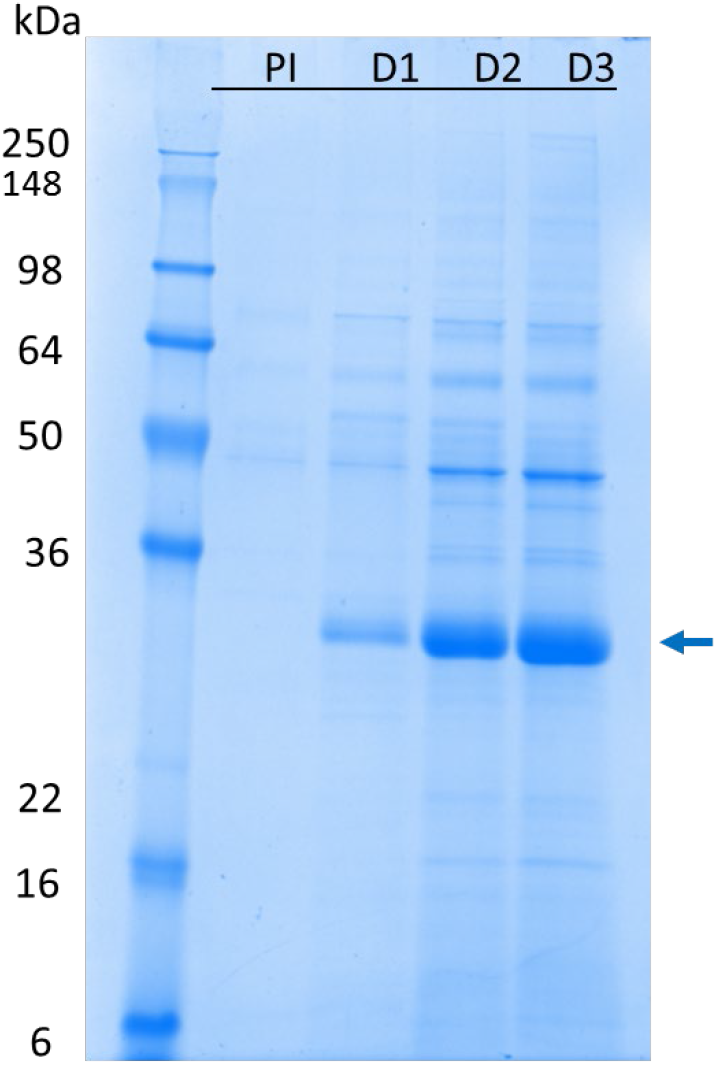
Time point SDS-PAGE analysis of pre- and post-induction fermentation samples of the lockdown process (Run 5). PI: pre-induction; D1, D2, D3: Day 1-3 after induction. The arrow shows RBD219-N1C1 in fermentation supernatant after induction.

### Three purification schemes

In parallel with the fermentation optimization, three different processes were performed to purify RBD219-N1C1 from the FS (Figure 1B). Process-1 was developed by adapting the purification method of our SARS-CoV-RBD219-N1 antigen that shares 70% homology with the SARS-CoV-2 RBD [8,4,27]. In this process, the target protein was captured by HIC using a butyl HP column with 1 M ammonium sulfate salt for the binding. After the HIC, 67% of the target protein was recovered and purity significantly improved from 85.6% in the FS to 97.6% (Figure 3A). The target protein was concentrated using Amicon centrifugal concentrators and further polished by SEC using a Superdex 75 column. The SEC elution pool was then diluted to 2 mg/mL for storage. Overall, the final yield of the target protein using Process-1 was 188.8 mg/L FS (Figure 3A), representing a recovery of 50% with a purity of 98.3%. This is similar to the overall recovery of 52% and the purity of 98.5% shown with the SARS-CoV-RBD219-N1 protein [4].

**Figure 3.**
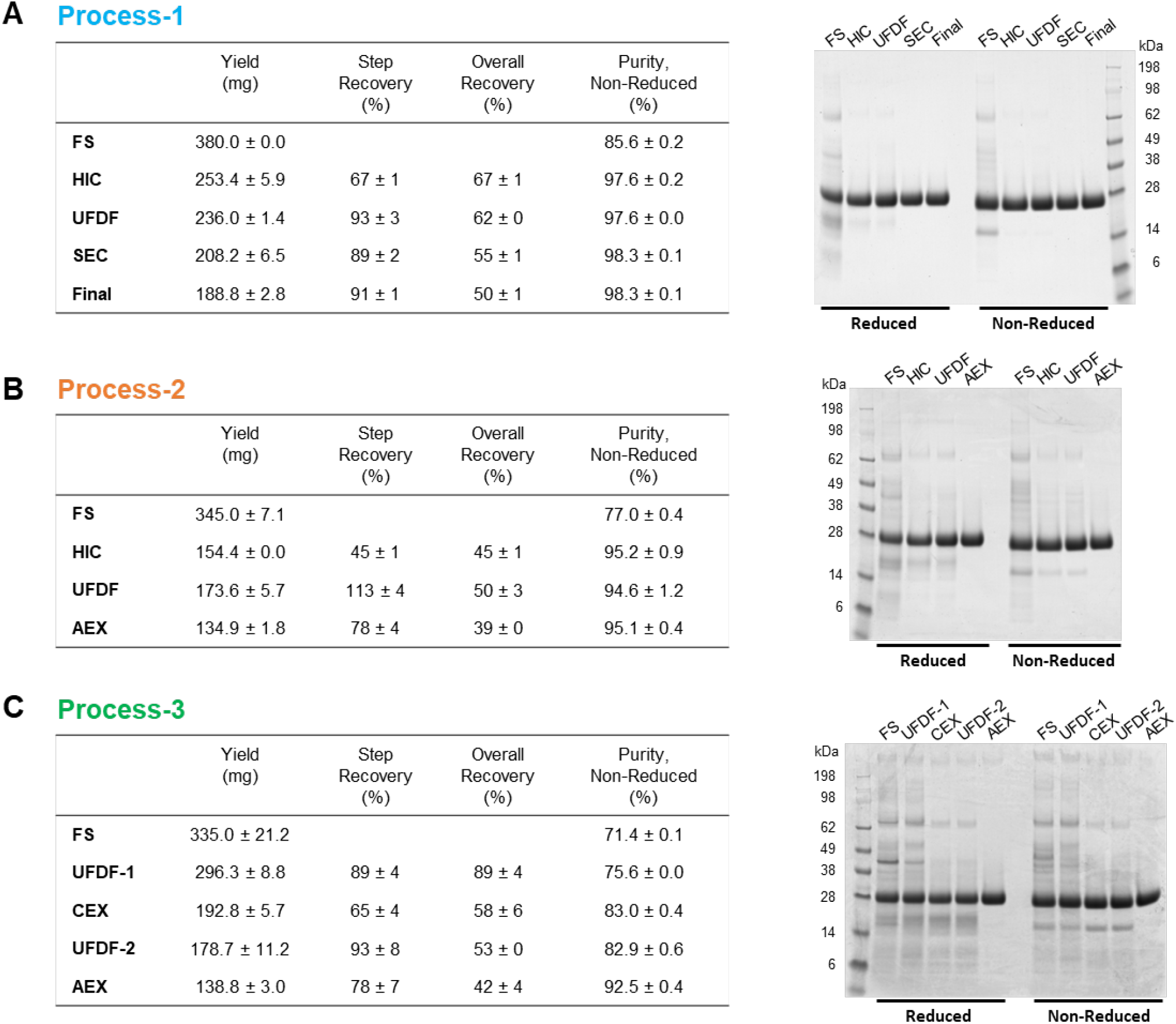
In-process samples comparison from three processes. (A-C) Yield, step recovery, overall recovery, and purity are shown as an average ± SD calculated from two independent gels are shown in table (left) and a representative gel stained with Coomassie Blue is shown (right) from Process-1 (A), Process-2 (B) and Process-3 (C). FS, fermentation supernatant; HIC, hydrophobic interaction chromatography; UFDF, Ultrafiltration, and diafiltration; SEC, size exclusion chromatography; AEX, anion exchange chromatography; CEX, cation exchange chromatography

Although Process-1 is sufficient and proven to produce proteins at high yield and purity at the laboratory scale, up to 10 L [4], there are considerations to be made with respect to scaling-up manufacture. Both HIC and SEC are costly steps due to their low binding and process capacities, requiring large resin volumes and long processing times. Therefore, we explored two alternative processes utilizing IEX, favored in the biopharmaceutical industry due to its low cost and high scalability.

In Process-2, the capture step was unchanged. After the UFDF step to concentrate and exchange buffer, the target protein was polished by a negative capture using AEX instead of SEC (Figure 1B). While the RBD219-N1C1 did not bind to the Q XL column, non-specific HCPs were bound to the column and removed effectively. The step recovery during AEX was 78%, which is lower than the 89% of the step recovery seen from SEC in Process-1. The final purity of the purified protein from Process-2 was 95.1%, which is lower than the 98.3% purity seen in Process-1 but still highly pure. However, the overall recovery in Process-2 was only 39%, much lower than the 50% for Process-1. This is due to the lower recovery during HIC, 45%, that lags the 67% recovery seen for the equivalent step in Process-1. This lower recovery may offer the opportunity for improvement, but overall, it is fair to conclude that AEX can successfully replace SEC for the polishing step.

In Process-3, we further optimized Process-2 to utilize CEX for the capture step instead of HIC. After the first UFDF step (UFDF-1) to concentrate and buffer exchange the FS, RBD219-N1C1 was captured using a CM FF column followed by a second UFDF step (UFDF – 2) and a polishing step using negative AEX capture (Figure 1B). The additional UFDF-1 step required prior to CEX increased processing time compared to Processes 1 and 2. Although the step recovery from CEX was 65%, very similar to the 67% seen after the HIC in Process-1, the purity was only 83% after CEX which is significantly lower than the purity (97.6% and 95.2% from Process-1 and −2, respectively) after HIC (Figure 3C). Purity was improved significantly by 9.5% after the polishing step, resulting in an overall purity of 92.5%, which is lower than the 98.3% and 95.1% seen in Processes 1 and 2, respectively.

To summarize, HIC showed a superior performance to remove non-specific host proteins and, hence, resulted in >95% purity after the capture step, which is even higher purity than 92.5% purity seen in the final protein product from Process-3. This favored HIC over CEX although its only drawback is the cost. HIC has no limitation on scale-up. On the other hand, both AEX and SEC showed very similar performance during the polishing step. However, while AEX is cost-effective chromatography with full scalability, SEC is expensive and has limitations in scale-up. This reasons us to favor Process-2 employing HIC and AXE for the capture and the polishing step, respectively. Before we urge to conclude that Process-2 is the best process to produce RBD219-N1C1, we characterized and compared the purified protein from Process-2 and two other processes for integrity, size estimation, impurity contents, and functionality.

### Characterization and size estimation of the purified proteins from three processes

The purified proteins from all processes were characterized for integrity by Western blot, SEC-HPLC, and DLS. When 5 μg of purified protein were analyzed by SDS-PAGE followed by Coomassie Blue staining, a single band was seen at ~28 kDa under reducing and ~25 kDa under non-reducing conditions (Figure 4A). Western blot analysis using a monoclonal antibody against SARS-CoV-2 Spike protein under a non-reducing SDS-PAGE indicating that the ~25 kDa band is indeed derived from the SARS-CoV-2 Spike protein (Figure 4B). An additional band at ~50 kDa was detected in the protein from Process-3, likely representing a dimer. Dimerization through free cysteine residues had also been reported for SARS-CoV-RBD219-N1, and therefore the free cysteine (C538) was mutated to alanine in RBD219-N1C1 [8]. Although RBD219-N1C1 theoretically lacks free cysteine residues, we observed some dimers during the fermentation that were removed during purification. Therefore, Process-3 appears to be less efficient at removing dimeric RBD219-N1C1 than the other processes.

**Figure 4.**
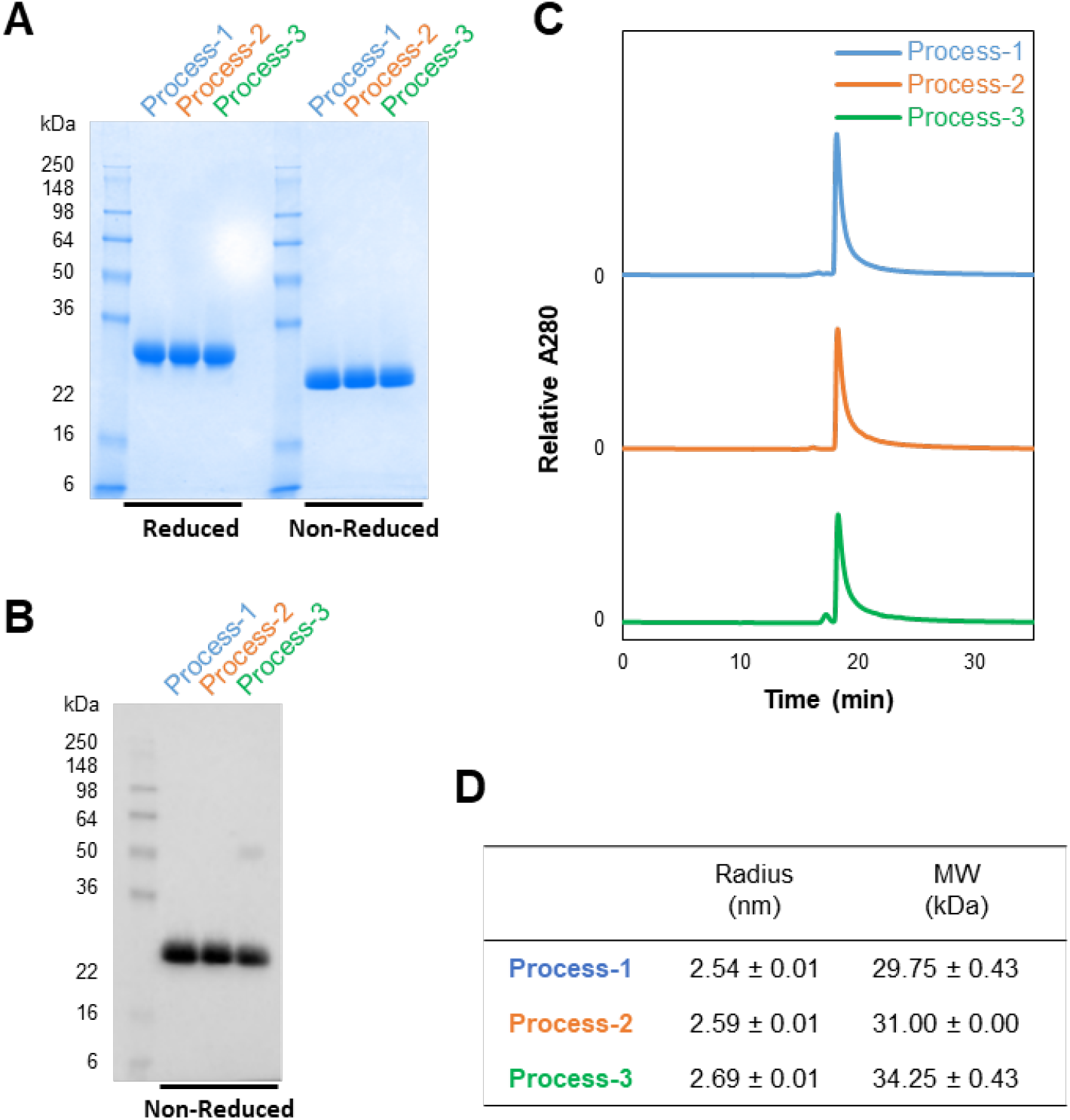
Characterization of purified RBD219-N1C1 proteins from three processes. (A-B) Purified proteins are analyzed by SDS-PAGE gel with Coomassie Blue stain (A) and Western blot with a monoclonal SARS-CoV-2 Spike antibody (B). (C-D) Size and aggregates evaluation by SEC-HPLC (C) and the radius and size in solution measured by Dynamic Light Scattering (D). (D) Averages ± SD are shown from four independent measurements.

SEC-HPLC with 50 μg of the purified protein preparations indicated that all three proteins were similar in size and had no aggregation. Only the purified protein from Process-3 showed an additional peak eluting ~1 min earlier, likely, as reported above, a dimer (Figure 4C). Finally, all three proteins were analyzed by DLS to estimate size and dispersity in solution. The estimated sizes of the purified proteins from each process were 29.75, 31.00, and 34.25 kDa, respectively (Figure 4D). As expected, the protein from Process-3 showed higher polydispersity than the other samples (Supplementary Figure 1).

### Impurity assessment in the purified proteins

*P. pastoris* HCP was assayed by Western blot and quantified by ELISA using a second-generation *P. pastoris* HCP detection kit. When 5 μg unpurified proteins (i.e. FS), as well as the purified proteins, were analyzed by SDS-PAGE followed by Coomassie Blue stain and Western blot, we saw that HCP had been effectively removed from all three processes (Figure 5A and B). The HCP content in the purified proteins was calculated as 95.9 ng, 6.8 ng, and 44.8 ng per mg of RBD219-N1C1 from Processes 1-3, respectively (Figure 5C). All these values were within acceptable limits, 1-100 ng/mg, for biopharmaceuticals [2,30].

**Figure 5.**
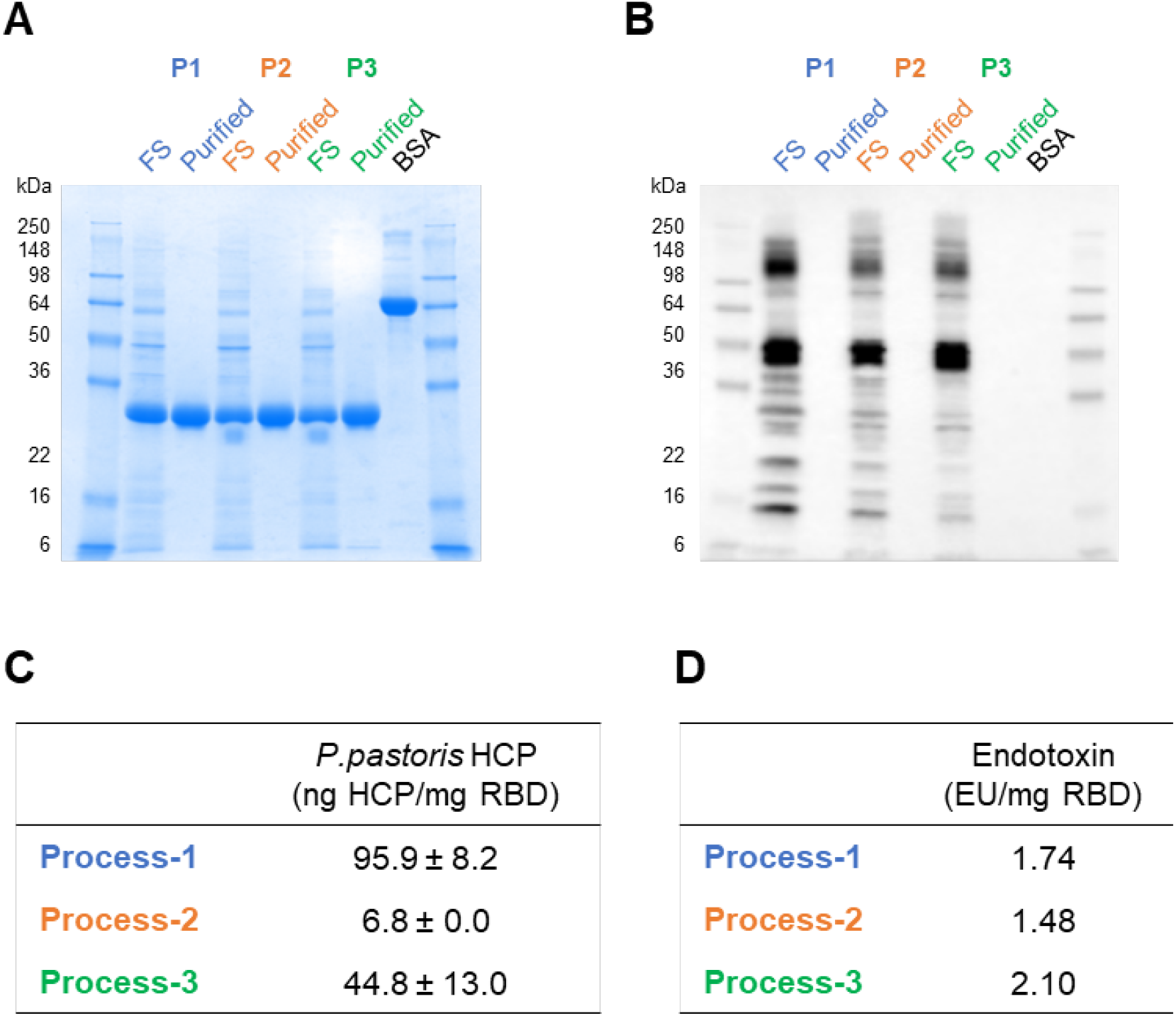
Impurity evaluation of the purified RBD219-N1C1 proteins from three processes. (A-B) Unpurified (FS) and purified RBD219-N1C1 in reduced SDS-PAGE with Coomassie Blue stain (A) and with Western blot using anti-*P. pastoris* HCP antibody (B). (C) Measured *P. pastoris* HCP content by quantitative ELISA and (D) endotoxin levels are shown.

Endotoxin levels measured in the purified proteins were 1.74, 1.48, and 2.10 EU per mg for the purified proteins from Processes 1-3, respectively (Figure 5D). These values are significantly lower than the maximum recommended endotoxin level for recombinant subunit vaccines, 20 EU/mL [3].

### Functionality assessment using ACE2 binding assay

To evaluate the functionality of the purified proteins from each process, the ability to bind to the human ACE2 receptor was tested *in vitro.* SARS-CoV-2 uses this human cell surface receptor for cell entry [14], and here the binding of each protein to ACE2 was quantified by ELISA. All proteins presented similar binding curves to ACE2, with calculated EC50-values (for 2 μg/mL purified protein) of 0.037, 0.033, and 0.038 μg/mL ACE2, respectively (Figure 6), suggesting that all three proteins were functionally equivalent.

**Figure 6.**
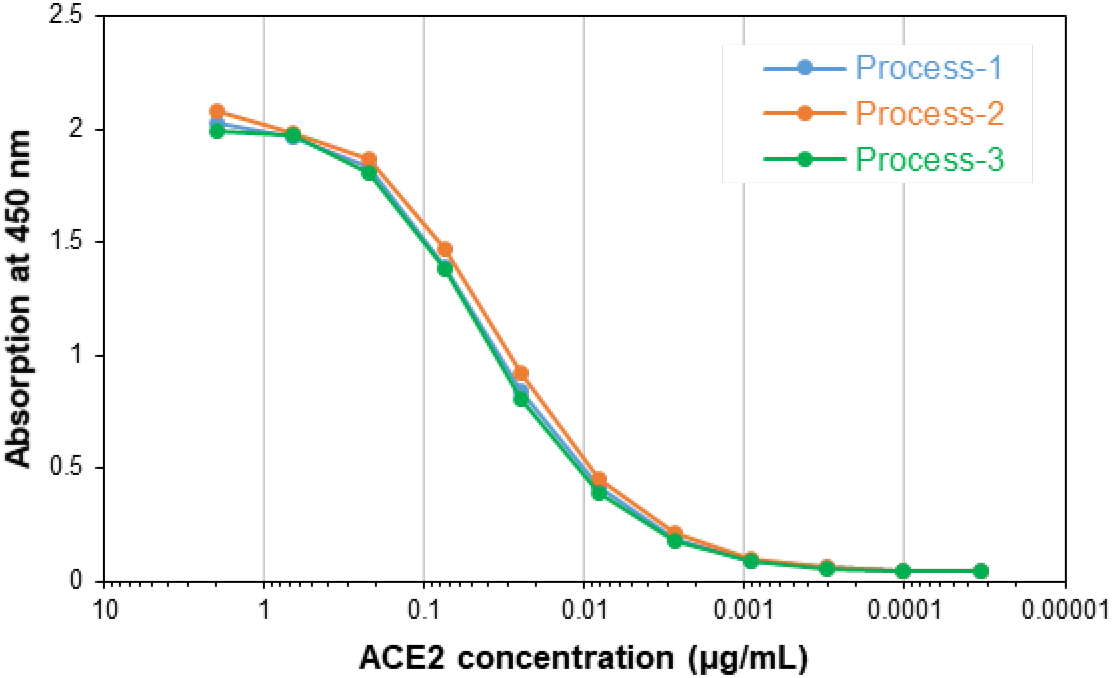
Binding ability of the purified RBD219-N1C1 from three processes to a recombinant human ACE2 receptor

## Discussion

We developed a process suitable for producing a recombinant protein COVID-19 vaccine antigen for clinical testing and transition to industrial manufacture. Fermentations were initially run at the 1 L (for fermentation conditions optimization) and then the 5 L scale (for downstream purification process development). When scaled to 5 L and conditions had only been modified for gas flow and agitation rate to maintain 30% dissolved oxygen, differences in the protein yield were observed. Four subsequent identical 5 L fermentation runs showed high reproducibility with a CV of 8.3%, further emphasizing robustness.

Based on our previous experience with SARS-CoV-RBD219-N1 (a prototype vaccine for SARS), a 1.6-to 2.5-fold yield increase was achieved when switching from basal-salt to low-salt medium during the glycerol batch phase in the fermentation process [5]. For SARS-CoV2-RBD219-N1C1, a 3.6-fold increase in yield suggests that the salt concentration was a significant factor. In basal-salt medium, the recombinant protein precipitates in the presence of magnesium and calcium phosphates as the pH is adjusted above 5.5. Low-salt medium also precipitates, though to a much lesser extent. The precipitate formation can have adverse effects on the fermentation process as it can lead to an unbalanced nutrient supply, cause cell disruption, and induce secreted proteins to form aggregates [29]. Similar findings had previously been observed with the production of a therapeutic Fc-fusion protein in the fermentation of *P. pastoris*. When salt supplements were added at induction, the protein yield decreased [23].

Purification optimization produced RBD219-N1C1 at high purity and yield, with a high recovery rate, suitable for scalability for manufacturing. Three purification methods (Processes 1-3) were tested and compared using 1 L FS from the identical fermentation runs for rapid development. Process-1 was adapted based on the previous purification method with SARS-CoV-RBD219-N1 with slight modification on ammonium salt concentration in HIC. Process-1 resulted in 98.3% purity with a 50% overall recovery rate, similar to the 98.5% purity and 52% overall recovery shown in SARS-CoV-RBD219-N1 purification (Figure 3A) [4]. Purity was dramatically increased to >97% after the HIC capture step (Figure 3A). Process-1 is suitable to produce the target protein at the laboratory scale but is limited in scale-up due to low binding and process capacities, as well as the long processing time leading to high cost for production. Therefore, two other processes were tested to replace costly HIC and SEC with CEX and AEX, respectively. For biopharmaceuticals, IEX has been favored in chromatography due to its robustness and full scalability [4].

While IEX was tested in the polishing step of Process-2, it was used for both capture and polishing steps in Process-3. In Process-2, AEX showed comparable step recovery, and purity increases to SEC in Process-1 (Figure 3A and B). However, a significant improvement in purity by AEX was shown in Process-3 after the capture step by CEX (Figure 3C) as the CEX elution pool showed only 83.0% purity. This suggests that AEX not only can successfully replace SEC but also can effectively remove non-specific host proteins. On the contrary, CEX showed comparable step recovery but a lower capability to remove host proteins during the capture step. The purity after the CEX-capture was only 83.0%, which is significantly lower than the purity after HIC-capture (97.6% and 95.2% seen in Processes 1 and 2, respectively) (Figure 3). Overall, Process-3 produced the least pure RBD219-N1C1 protein among the three processes tested.

Before choosing the best process for purification of RBD219-N1C1, the purified proteins were characterized for their quality based on size, specificity, and impurity. The integrity assessment of the purified proteins was performed by SDS-PAGE. Coomassie-stained gels showed a single band at ~25 kDa that was recognized by a SARS-CoV-2 spike protein-specific antibody (Figure 4A and B). In addition, for Process-3, the western blot indicated the presence of an additional band speculated to be a dimer (Figure 4B); this same product was also seen by SEC-HPLC (Figure 4C). Although no difference in size was seen among the purified proteins from the three processes by SDS-PAGE (Figure 4A), the size under native conditions, estimated by DLS showed differences. The sizes in solution were 29.75, 31.00, and 34.25 kDa for the products from Processes 1-3, respectively. The purified protein from Process-3 appeared larger estimated size suggesting the presence of additional molecules in the preparation (Figure 4D and Supplementary Figure 1). Next, impurities such as *P. pastoris* HCPs and endotoxin levels were analyzed and compared. While all purified proteins showed no detectable HCPs by western blot with anti-*P. pastoris* antibodies (Figure 5B), when measured by ELISA different HCP content levels were observed. Process-2 showed the lowest HCP content (6.8±0.0 ng) per mg of purified protein while Process-1 showed the highest HCP content (95.9±8.2 ng) and Process-3 showed 44.8±13.0 ng (Figure 5C). The higher HCP content found in the purified protein from Process-1 was likely due to the presence of HCP with a similar size of RBD219-N1C1, which further suggested that SEC might not be an ideal purification step. No significant difference in endotoxin level was measured in the purified protein from three processes (Figure 5D), albeit all protein preparations contained less than the maximally allowed endotoxin levels. Finally, the functionality of the purified proteins from three processes tested by in vitro ACE2 binding assay showed that all three proteins showed similar binding to recombinant human ACE2 receptor (Figure 6).

In summary, after comparing yield, purity, and recovery after each purification, we conclude that HIC for capture due to its superior capability to remove non-specific host proteins and produce protein with >95% purity, and AEX for polishing due to its low cost and full scalability (Process-2) are best suited to produce RBD219-N1C1. In addition, comparison for the integrity, dimer content, HCP contents, and endotoxin level in purified protein supported the Process-2 generates quality proteins similar to Process-1 but significantly better than Process-3 and, hence, is a more ideal process for upscaling.

*P. pastoris* is widely used to produce recombinant proteins for clinical and commercial use. The *P. pastoris* system is licensed to more than 300 companies in the biotechnology, pharmaceutical, vaccine, animal health, and food industries, and more than 70 therapeutic and industrial products are approved by stringent regulatory bodies including human insulin, Hep B vaccine, cytokines, and hormones [26]. *P. pastoris* offers high growth rates, high cell densities, and high protein yield using simple and inexpensive fermentation media. Fermentation conditions are highly scalable due to the robust nature of *P. pastoris*, and the manufacturing times are short. With such an effective production platform and the availability of manufacturing facilities including vaccine manufacturers from the developing countries network, we can produce this COVID vaccine candidate at a low cost to meet the urgent global needs. The production technology of RBD219-N1C1 was transferred to Biological E. Limited, an India-based vaccine and pharmaceutical company, and a Phase I/II clinical trial was initiated in November 2020 in India [11,10].

## Contribution

JL, ZL, WC, PG, US, PH and MB conceived and designed research. JL, ZL, WC, RK, RA, and JR conducted experiments and analyzed data. Everybody contributed to discussing the results and writing the manuscript. All authors read and approved the manuscript.

## Declaration of interest statement

No potential conflicts of interest were disclosed.

## Funding

This work was supported by the Robert J. Kleberg Jr. and Helen C. Kleberg Foundation; Fifth Generation, Inc. (Tito’s Handmade Vodka); JPB Foundation, NIH-NIAID (AI14087201); and Texas Children’s Hospital Center for Vaccine Development Intramural Funds. We also would like to thank PATH Center for Vaccine Innovation and Access (Seattle, WA, USA) for their guidance as well as technical and intellectual support.

---

**Supplementary Figure 1.**
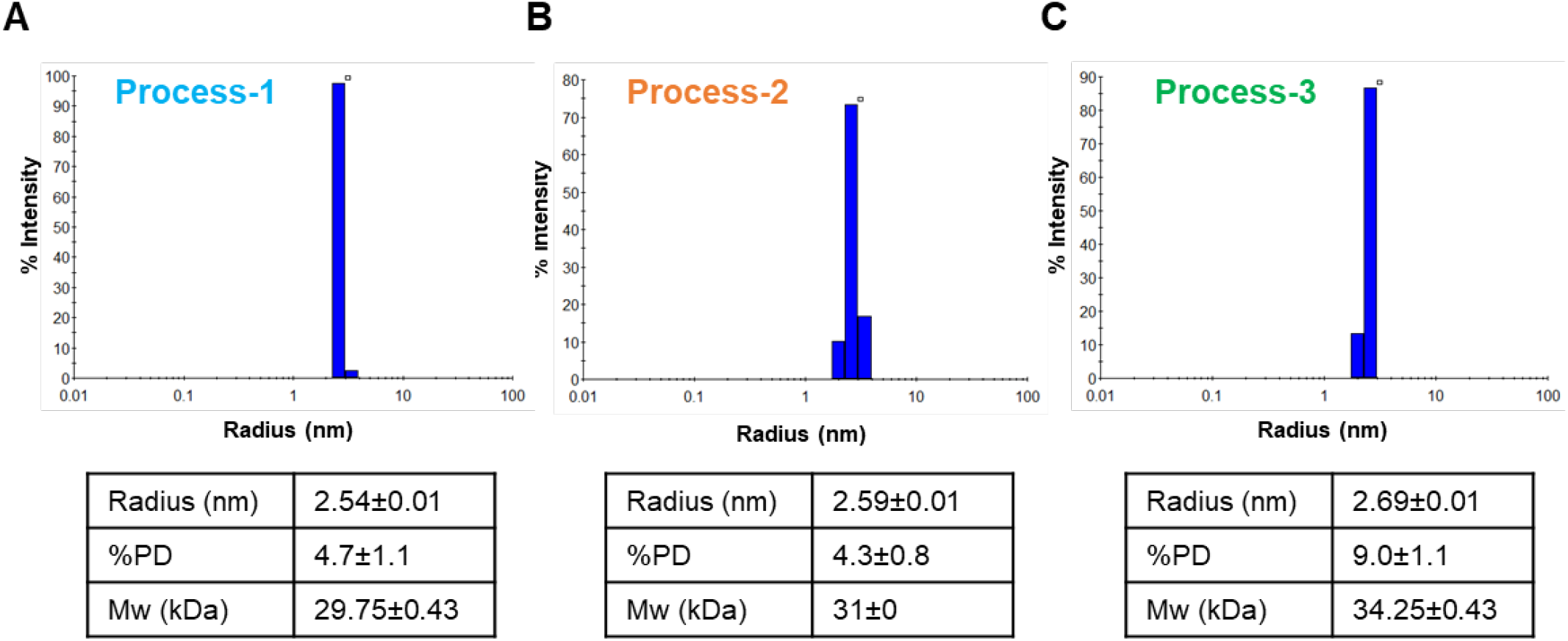
Dynamic light scattering results of the purified proteins by Process-1 (A), Process-2 (B), and Process-3 (C). Measured Stokes radii, polydispersity (PD), and molecular weight (Mw) are shown as an average ± SD from four independent measurements.

## Notes

### Competing Interest Statement

The authors have declared no competing interest.

